# Development of a Customizable Pacing Protocol to Induce Persistent Atrial Fibrillation in Swine

**DOI:** 10.64898/2025.12.01.691684

**Authors:** Fox Bravo, Jacob Ref, Jesse Riemenschneider, Eli Lefkowitz, Pouria Mostafizi, Alan Salciccioli, Sophia DiPonio, Adrian Grijalva, Daniel Benson, Allison S. Tulino, Mary Kay Pierce, Steven Goldman, Talal Moukabary

## Abstract

**Introduction:** Persistent atrial fibrillation (AFib) research is dependent on large animal models to understand pathogenesis and test new treatments and therapeutic techniques. While various methods have been established to induce persistent AFib in large animals, including electrical, surgical, pharmacological, and genetic approaches, each has distinct limitations. The most recent being the device industry no longer providing off-target rapid atrial pacing programs for use in animals. This challenge requires the development of a new accessible model of inducing atrial fibrillation in large animals.

**Methods Used:** We developed a wireless pacing system using a Raspberry Pi Pico W microcontroller (Pico) programmed for variable pacing frequencies. The device is powered by a subcutaneously implanted 9V battery. The Pico’s built-in Wi-Fi capabilities enable remote connection and real-time adjustment of pacing frequency output.

This protocol describes a prospective study in which swine will undergo chronic right atrial pacing for 3-4 weeks to induce persistent AFib. We tested our device in a domestic swine. Vascular access was established through the left jugular vein and an active fixation pacing lead (Medtronic 5076) was implanted under fluoroscopic guidance in the right atrium. Proper lead positioning and pacing function were confirmed through electrocardiographic monitoring of both atrial and ventricular capture.

**Preliminary Results:** Oscilloscope testing demonstrated frequency and voltage output concordant with the programmed frequencies while real-time adjustments were made through the Wi-Fi interface. A cardiac pacing wire was placed in the right ventricle then relocated to the right atrium and successful pacing with capture was verified using an electrocardiogram.

**Conclusions:** This protocol will provide a system capable of capturing atrial tissue with confirmed wireless power transfer capabilities that has minimal tissue heating and is physiologically safe. Combined with our literature review findings that electrical atrial pacing methods are most effective for inducing persistent AFib, our device provides researchers with the potential for an improved tool for creating large animal models of persistent AFib.

## Introduction

Atrial fibrillation (AFib) is the most common persistent arrythmia in adults in the United States. Atrial fibrillation currently affects between 3-6 million people in the U.S. with this number expected to rise with the growing population (1, 2). Given the significant disease burden, developing robust large animal models of AFib provides ability to test novel pharmacologic and interventional therapies.

The pathophysiology of AFib is well described in the literature with knowledge about the origin of ectopic beats and the genetic and structural changes that occur. The electrical remodeling in AFib is driven by the changes in ion channels, notably the downregulation of L-type calcium channels (3). This downregulation in calcium channels can cause the generation of reentrant circuits which increases the susceptibility to AFib. Atrial fibrillation induces fibrotic changes driven by fibroblast activation leading to excessive extracellular matrix deposition which disrupt the electrical conduction (4). The autonomic nervous system also influences AFib susceptibility with sympathetic innervation enhancing automaticity and early afterdepolarization, while parasympathetics shorten the atrial refractory period which increases risk for reentrant activity (5). Recently, genetic factors such as variants of the PITX2 gene have been associated with increased AFib risk (6). Furthermore, microRNA dysregulation and epigenetic modifications also contribute to AFib pathogenesis through fibrosis and changes in ion channel expression, both of which are potential targets for new therapeutics (7). These underlying changes lead to ectopic activity, particularly in the pulmonary veins, which are understood to be susceptible because these veins have myocardial sleeves capable of spontaneous depolarization (8).

Researchers have established various animal models of AFib that can induce either transient or chronic sustained arrhythmia. Swine have served as the mainstay in translational AFib research due to their similar cardiac physiology and ability to use catheter ablation-based techniques (9, 10). The model we review in this article aims to reproduce the previously described pathophysiology to induce AFib in swine.

### Electrical Models of Atrial Fibrillation

Many methods of inducing AFib in swine have been created (Table 1). Electrical stimulation is the most reliable and reproducible method, traditionally split into two subcategories: atrial tachypacing and burst pacing. Atrial tachypacing involves persistent pacing of the right atrium at a fixed rapid pace above the intrinsic atrial rate, inducing electrophysiologic and structural changes. Burst pacing delivers high-frequency stimuli to the atrium at a rate shorter than the refractory period, aiming to induce AFib by creating premature depolarizations and reentrant circuits.

**Table 1.**
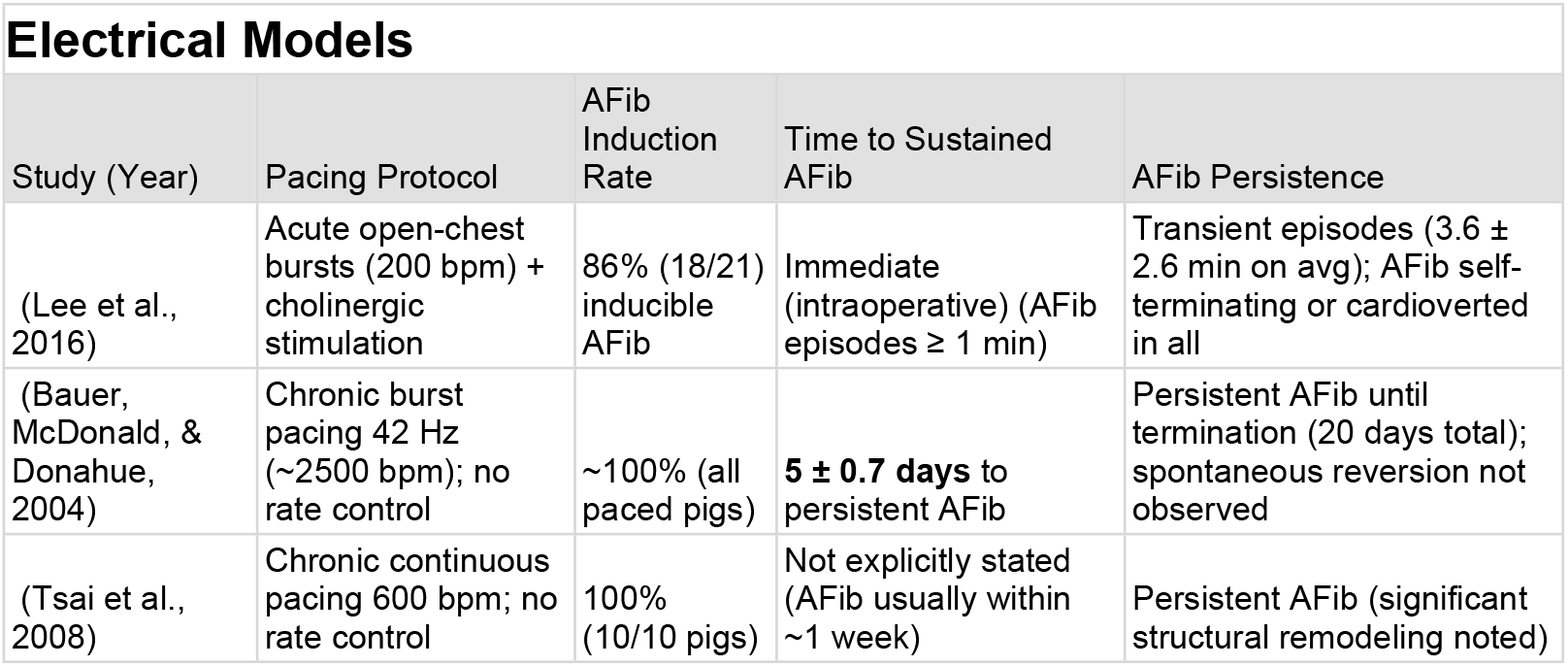
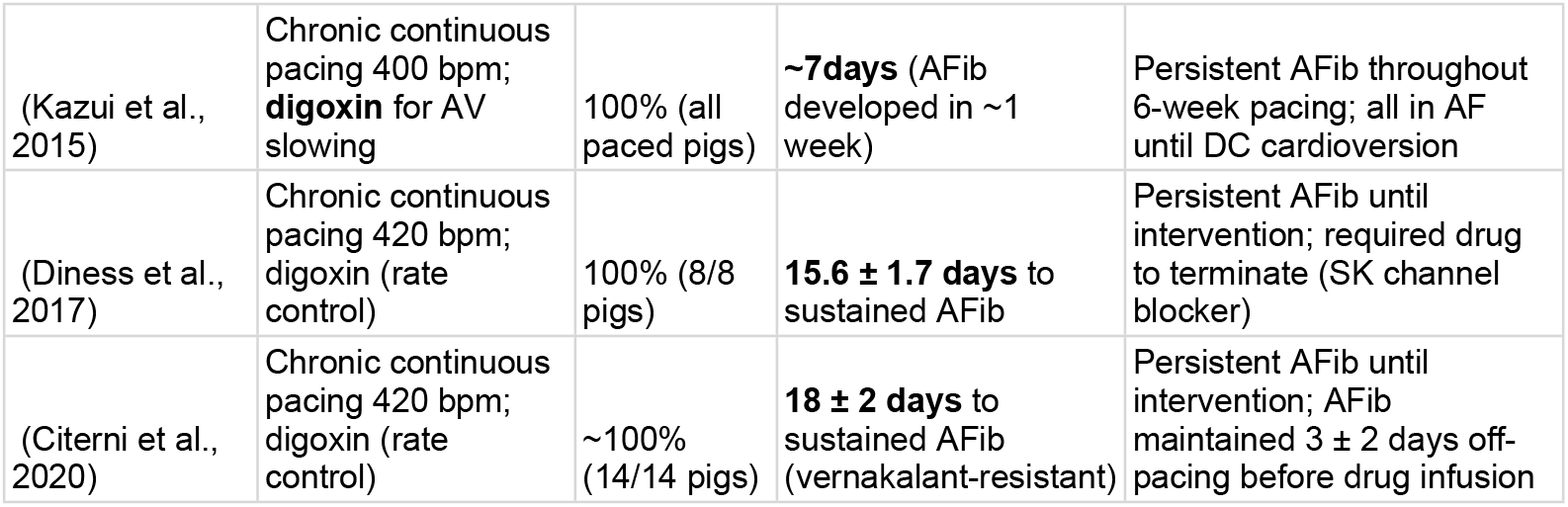
Outlines electrical models of inducing atrial fibrillation in swine. This table includes the study (a) author/ year, (b) protocol used, (c) number of swine that converted from sinus rhythm, (d) time required to induce AFib, (e) how long AFib persisted following pacing.

| Study (Year) | Pacing Protocol | AFib Induction Rate | Time to Sustained AFib | AFib Persistence |
| --- | --- | --- | --- | --- |
| (Lee et al., 2016) | Acute open-chest bursts (200 bpm) + cholinergic stimulation | 86% (18/21) inducible AFib | Immediate (intraoperative) (AFib episodes $\geq 1$ min) | Transient episodes ($3.6 \pm 2.6$ min on avg); AFib self-terminating or cardioverted in all |
| (Bauer, McDonald, & Donahue, 2004) | Chronic burst pacing 42 Hz ($\sim 2500$ bpm); no rate control | $\sim 100\%$ (all paced pigs) | **$5 \pm 0.7$ days** to persistent AFib | Persistent AFib until termination (20 days total); spontaneous reversion not observed |
| (Tsai et al., 2008) | Chronic continuous pacing 600 bpm; no rate control | 100% (10/10 pigs) | Not explicitly stated (AFib usually within $\sim 1$ week) | Persistent AFib (significant structural remodeling noted) |
| (Kazui et al., 2015) | Chronic continuous pacing 400 bpm; **digoxin** for AV slowing | 100% (all paced pigs) | **~7days** (AFib developed in ~1 week) | Persistent AFib throughout 6-week pacing; all in AF until DC cardioversion |
| (Diness et al., 2017) | Chronic continuous pacing 420 bpm; digoxin (rate control) | 100% (8/8 pigs) | **15.6 ± 1.7 days** to sustained AFib | Persistent AFib until intervention; required drug to terminate (SK channel blocker) |
| (Citerni et al., 2020) | Chronic continuous pacing 420 bpm; digoxin (rate control) | ~100% (14/14 pigs) | **18 ± 2 days** to sustained AFib (vernakalant-resistant) | Persistent AFib until intervention; AFib maintained 3 ± 2 days off-pacing before drug infusion |

**Table 2.** Summarizes genetic manipulations that have been used to prolong onset of atrial fibrillation in swine. The table includes (a) author/ year, (b) genetic modification used, (c) days to AFib onset in genetically modified swine and controls (unmodified pigs), (d) success rate of AFib induction, (e) how long AFib persisted following pacing. For context, control pigs in these studies nearly always developed persistent AFib within ∼5–7 days (100% incidence) under the same pacing protocols.

| Genetic Models |  |  |  |  |
| --- | --- | --- | --- | --- |
| Study (Year) | Genetic Modification | Days to AFib Onset | AFib Induction Success | AFib Duration Once Induced |
| (Soucek et al., 2012) | Adenoviral **KCNH2-G628S** | **12 ± 2.1 days** (treated) vs ~6.2 ± 1.3 days in controls | ~67% (some pigs never developed AFib in 14 days) vs **100%** in controls | **Persistent AFib** in those that did convert (none in 33% of treated pigs) |
| (Bikou et al., 2011) | Adenoviral **Connexin-43** (Cx43) gene transfer (gap junction up regulation) | **>14 days** (no AFib during 2- week pacing) vs $\sim 7.4 \pm 0.5$ days in controls | **0%** (0/5 pigs) developed AFib in 14 days vs **100%** of controls | **No sustained AFib** observed in treated pigs (controls had persistent AFib) |
| (Igarashi et al., 2012) | Adenoviral **Connexin-40 + Connexin-43** co-transfer (improved coupling) | **>7 days** (no continuous AFib during one-week study) vs $\sim 5.8 \pm 0.6$ days in controls | **0%** with continuous AF by 7 days vs **100%** in controls | **No persistent AFib** in treated pigs over 7 days |
| (Liu et al., 2017) | Adenoviral **KCNH2-G628S** (Same as Souček, tested in post-operative AF model) | **N/A** (evaluated by % time in sinus rhythm during 14 days pacing) | Treated pigs in sinus **59%** of time vs **14%** in controls ($p=0.009$) | **Fewer/shorter AFib episodes** with gene therapy (significantly reduced AFib burden) |
| (Park et al., 2015) | **SCN5A E558X/+** mutation (germ line transgenic pig; Na<sup>+</sup> channel loss-of-function) | No spontaneous AFib observed over **>2 years** (conduction disease + VF propensity noted) | AFib **0%** spontaneously (model predisposed mainly to ventricular arrhythmias) | N/A (no AFib; exhibited **ventricular fibrillation** on induction) |

**Table 3.**
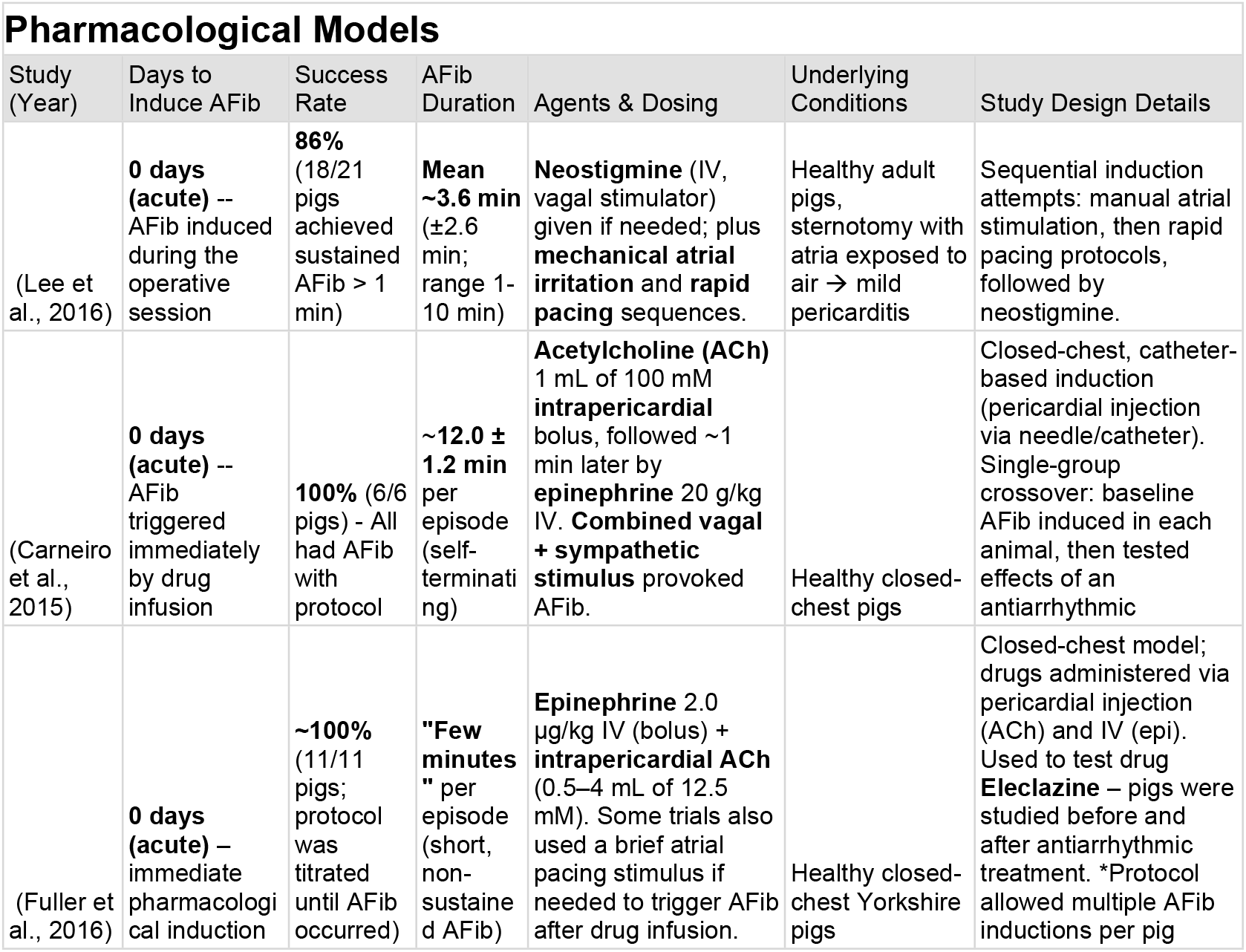
Outlines pharmacologic methods of inducing atrial fibrillation in swine. The table includes (a) author/ year, (b) time required to induce AFib, (c) success rate of AFib induction, (d) how long AFib persisted following pacing, (e) type of pharmacologic/s used, (f) health of the swine, (g) protocol used to induce AFib pharmacologically.

| Study (Year) | Days to Induce AFib | Success Rate | AFib Duration | Agents & Dosing | Underlying Conditions | Study Design Details |
| --- | --- | --- | --- | --- | --- | --- |
| (Lee et al., 2016) | **0 days (acute)** -- AFib induced during the operative session | **86%** (18/21 pigs achieved sustained AFib > 1 min) | **Mean ~3.6 min** ($\pm 2.6$ min; range 1-10 min) | **Neostigmine** (IV, vagal stimulator) given if needed; plus **mechanical atrial irritation** and **rapid pacing** sequences. | Healthy adult pigs, sternotomy with atria exposed to air → mild pericarditis | Sequential induction attempts: manual atrial stimulation, then rapid pacing protocols, followed by neostigmine. |
| (Carneiro et al., 2015) | **0 days (acute)** -- AFib triggered immediately by drug infusion | **100%** (6/6 pigs) - All had AFib with protocol | **$\sim 12.0 \pm 1.2$ min** per episode (self-terminating) | **Acetylcholine (ACh)** 1 mL of 100 mM **intrapericardial** bolus, followed $\sim 1$ min later by **epinephrine** 20 g/kg IV. **Combined vagal + sympathetic stimulus** provoked AFib. | Healthy closed-chest pigs | Closed-chest, catheter-based induction (pericardial injection via needle/catheter). Single-group crossover: baseline AFib induced in each animal, then tested effects of an antiarrhythmic |
| (Fuller et al., 2016) | **0 days (acute)** -- immediate pharmacological induction | **$\sim 100\%$** (11/11 pigs; protocol was titrated until AFib occurred) | **"Few minutes"** per episode (short, non-sustained AFib) | **Epinephrine** 2.0 μg/kg IV (bolus) + **intrapericardial ACh** (0.5–4 mL of 12.5 mM). Some trials also used a brief atrial pacing stimulus if needed to trigger AFib after drug infusion. | Healthy closed-chest Yorkshire pigs | Closed-chest model; drugs administered via pericardial injection (ACh) and IV (epi). Used to test drug **Eleclazine** – pigs were studied before and after antiarrhythmic treatment. *Protocol allowed multiple AFib inductions per pig |

**Table 4.** Presents surgical models of inducing atrial fibrillation in swine. The table includes (a) author/ year, (b) surgical method used to induce AFib (c) time required to induce AFib, (d) number of pigs successfully converted, (e) duration of AFib following induction.

| **Surgical Models** |  |  |  |  |
| --- | --- | --- | --- | --- |
| Study (Year) | Induction Method | Days to AFib Onset | Success Rate | AFib Duration |
| (Lee et al., 2016) | Open-chest acute induction (manual atrial stimulation + pacing + neostigmine) | 0 days (immediate, intra-op) | 86% (18/21 pigs) | ~3.6 ± 2.6 minutes per episode (sustained ≥1 min) |
| (Lee et al., 2023) | Sterile pericarditis (post-op inflammation) + burst pacing | 1–3 days post-surgery (POAF induction window) | 43% of attempts (3 of 7 pigs) | ≥5 minutes per AFib episode (sustained POAF definition) |

Acute induction was performed in 21 pigs that underwent open-chest pacing with stepwise protocol aimed to induce short-term AFib episodes during surgical procedures (11). Chronic high-rate pacing with uncontrolled ventricular rate was performed by pacing at extremely high atrial rates without AV nodal rate control (12, 13). These protocols resulted in tachycardia-induced cardiomyopathy and heart failure due to very high ventricular rates. More recent methods used chronic high-rate pacing with controlled ventricular rate used atrial pacing at ∼400-420 bpm combined with AV nodal blocking agents to prevent excessive ventricular response (14, 15). Similar protocols were used by Kazui et al., 2015 (16).

### AFib Induction Success Rates

AFib induction success rates were uniformly high across pacing studies. Nearly all pigs subjected to adequate tachypacing developed sustained AFib. For instance in one study using atrial tachypacing persistent AFib was achieved in all paced pigs (12) with similar results in subsequent studies (8/8 pigs (15), 10/10 pigs (17), and 14/14 pigs (14)).

The time required to induce chronic AFib varied with pacing protocol. In rapid/uncontrolled ventricular response groups, Bauer’s study reported persistent AFib within 5.0 ± 0.7 days on average (12). When ventricular rate was pharmacologically controlled, AFib induction took longer, achieving sustained AFib in ∼17–19 days (18 ± 2 days (14) and 15.6 ± 1.7 days (15) on average). Rates ≥300 bpm were necessary to produce electrophysiological remodeling for AFib (18).

### Meta-analytic Findings

The pooled mean time to chronic AFib (persistent AFib >7 days) across all chronic studies was approximately 12–14 days of tachypacing. However, heterogeneity was high (Q-statistic p<0.001) due to protocol differences. Subgroup analysis underscores that pacing with uncontrolled ventricular response (VR) significantly shortens AFib induction time (5–7 days) compared to pacing with VR control (∼18 days, p<0.01). In practical terms, pacing at ≥300–400 bpm is required to induce AFib in pigs (18) but whether AFib manifests in days vs. weeks depends on the presence of rapid ventricular activation and resulting cardiomyopathy.

In summary, atrial tachypacing in pigs reproducibly induces persistent AFib. All reviewed protocols achieved high induction success of AFib, with differences in timing and heart failure development linked to pacing strategy. Statistically, protocols with an uncontrolled ventricular response had significantly shorter AFib induction times (by ∼12 days on average) at the cost of heart failure, whereas controlled-response protocols preserved ventricular function but required longer pacing to reach the persistent AFib stage. Researchers can thus tailor the model to their needs: a fast AFib + HF model (≈1 week) or a slower lone-AFib model (≈3 weeks).

### Genetic Models of Studying Atrial Fibrillation

Swine do not develop persistent AFib naturally; however, researchers have engineered genetic models to evaluate changes in AFib inducibility. These include ion channel mutations (19) and

(20) and gap-junction enhancements (connexin-43 or -40 gene transfer) (21, 22). All studies mentioned here consistently show that genetic modifications make persistent AFib more difficult to induce in pigs.

Introducing a dominant-negative KCNH2 (potassium voltage gated channel) mutation roughly doubled the time to induce persistent AFib (20). Similarly, in a post-operative AFib model, KCNH2-G628S gene therapy kept pigs in sinus rhythm ∼59% of the time vs only 14% in controls (19). Gap-junction enhancements had an even more striking effect; 0% of Cx43-treated pigs converted into AFib over 14 days, whereas all control pigs did by ∼7 days (22). A similar adenoviral study found that none of the connexin-treated pigs had continuous AFib during one week of rapid pacing (21).

To date, there is only one germline genetically engineered pig model relevant to arrhythmia, *SCN5A*^*E558X/+*^ pig (23). This model exhibited atrioventricular conduction block and propensity for ventricular fibrillation but did not demonstrate spontaneous atrial fibrillation.

In summary, the collected evidence demonstrates that these genetic pig models require more stimulation to initiate AFib and succeed less often in sustaining AFib compared to unmodified pigs. These findings not only validate the role of specific genes (ion channels, gap junctions) in AFib pathophysiology but also highlight promising genetic strategies for AFib prevention.

### Pharmacologic Models of Atrial Fibrillation

Pharmacological methods to induce AFib in pigs typically used two main approaches: cholinergic stimulation often combined with sympathetic stimulation, and direct autonomic agonist combinations. All studies achieved AFib acutely, often within minutes of drug administration (11, 24, 25).

Combined autonomic stimulation was very reliable – Carneiro’s team reported a 100% success rate using acetylcholine plus epinephrine. Another study induced controlled atrial ischemia, hypothesizing that ischemia plus adrenergic stress would increase AFib vulnerability (26). To study comorbidities that are commonly associated with AFib researchers have added hypertension via chronic deoxycorticosterone treatment to a pacing model (27).

All studies reported the ability to induce AFib pharmacologically, though AFib episodes were typically of short duration (on the order of minutes) and not sustained long-term. The duration depended on the method of AFib induction: higher adrenergic drive tended to prolong the duration of fibrillation. This trend highlights a limitation that while pharmacological triggers can initiate AFib, maintaining long AFib in pigs often still requires structural remodeling or continuous pacing (which were outside the scope of these acute studies). The studies also demonstrate that this method provides a useful platform for testing acute interventions: for example, both a novel sodium channel blocker and localized amiodarone therapy were shown to suppress induced AFib, suggesting the model’s relevance for evaluating pharmacotherapies.

### Surgical Models of Atrial Fibrillation

Surgical models induce chronic AFib by creating underlying structural heart disease. Approaches to induce AFib surgically include mitral regurgitation, hypertension and heart failure models, and inflammatory models. Investigators have created severe mitral regurgitation (MR) by surgically cutting the mitral chordae tendineae or damaging a papillary muscle (28). However, unless pacing or other triggers are added, MR alone typically leads to paroxysmal AFib rather than continuous AFib (29).

Post-myocardial infarction atrial arrhythmias can be studied by occluding coronary arteries (30). Studies using surgical techniques include open-chest atrial stimulation and pericarditis models (11, 31). The inflammatory substrate led to AFib episodes in ∼43% of attempts, consistent with the ∼40–50% success post operative AFib (POAF) induction rate seen in canine models (32).

## Methods

This protocol describes a prospective, controlled animal study designed to validate a novel, customizable wireless pacing system for inducing persistent atrial fibrillation in swine through chronic rapid atrial pacing. The study is conducted at the University of Arizona, Sarver Heart Center in AAALAC-accredited facilities equipped with appropriate surgical suites, fluoroscopy capabilities, and post-operative monitoring equipment. The primary aim is to demonstrate successful induction of persistent AFib (defined as continuous AFib lasting ≥7 days) using our custom device, while characterizing the associated electrophysiologic and structural remodeling.

### Animal Subjects and Ethical Compliance

One domestic swine aged 9-12 months old was used for device validation. This study is carried out according to the American Physiological Society and NIH Guidelines and was approved by the University of Arizona Institutional Animal Use and Care Committee (number 17-259, approved 07/05/2023).

The further subjects in this study will utilize mini-swine aged 9-15 months and weighing 60-80 kg at study initiation. Based on prior electrical pacing studies demonstrating approximately 100% AFib induction success rates, we calculated that n=6 mini-swine in the paced group provides greater than 80% power to detect successful AFib induction in at least 5 of 6 animals (83% success rate, 95% confidence interval: 44-97%). Inclusion criteria require normal baseline electrocardiogram (sinus rhythm), normal echocardiographic parameters (ejection fraction >50%), and absence of pre-existing cardiac arrhythmias or structural heart disease. Animals are excluded if they develop active infection, experience surgical complications preventing successful lead placement, or fail to achieve consistent pacing capture during implantation.

Animals will be randomly assigned to paced or control groups using computer-generated random numbers, and investigators performing terminal electrophysiology studies and histological analyses will be blinded to group assignment.

### Anesthetic Protocol and Surgical Procedures

A domestic swine weighing 70 kg was pre-anesthetized, intubated, and mechanically ventilated during the duration of the procedure. Standard aseptic surgical techniques were employed for all interventions. Vascular access was established through surgical cutdown technique targeting the jugular vein and carotid artery, with preemptive loose ligation applied to minimize intraoperative bleeding. The methods are further described in **Supplement 1**.

### Hardware Components and Materials

The pacing device was constructed using the following components: Raspberry Pi Pico W microcontroller with pre-soldered headers, universal printed circuit board (PCB), 7805 voltage regulator for power management, custom atrial pacing wire connection ports, medical-grade atrial pacing leads, 9V battery with adapter, and standard breadboard jumper wires for internal connections.

### Circuit Assembly and Configuration

The device assembly followed a standardized protocol. Headers were soldered to the Raspberry Pi Pico W to enable secure connections. Custom atrial pacing wire ports were attached to the microcontroller headers and positioned at the top right corner of the PCB with ports oriented away from the main processing unit (Figure 1). The electrode port for the distal pacing wire was connected to GPIO pin 11 (physical pin 15), while the cathode port was connected to ground pin 18 (GND 18).

**Figure 1.**
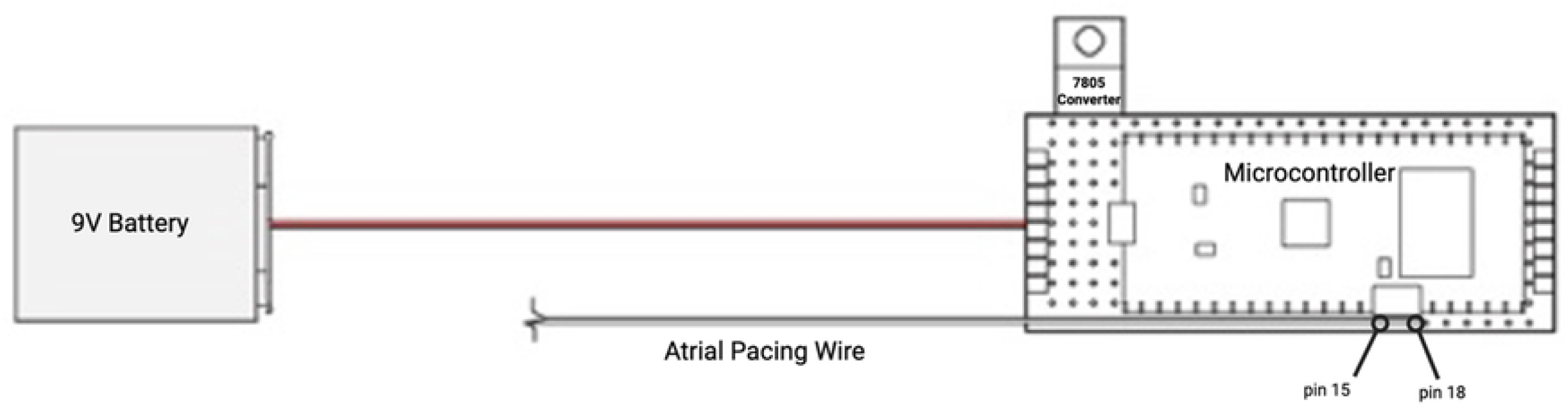
Assembly of the Raspberry Pi Pico W and it’s components: 9V battery, atrial pacing wire, 7805 voltage converter, and pins that the atrial pacing wire attach.

Power regulation was achieved through integration of a 7805 voltage regulator, with solder bridges connecting the regulator output to VSYS and the regulator ground to the nearest available ground connection (GND 3). The 9V battery adapter was integrated with solder bridges connecting the positive terminal to the voltage regulator input and the negative terminal to the regulator ground. Functional testing was performed following each assembly step.

### Software Implementation and Wireless Control

Custom firmware was developed and compiled into Universal Flash Format (UF2) files for deployment to the Raspberry Pi Pico W (**Supplement 2**). Firmware installation required connecting the device to a computer while holding the BOOTSEL button, followed by direct file transfer to the mounted device.

Wireless connectivity was implemented using the integrated Wi-Fi capabilities of the Raspberry Pi Pico W. Device configuration was managed through Visual Studio Code with the serial monitor extension. Wi-Fi network configuration required temporary hardware connection (connecting a designated wire to pin 1 for 8 seconds) followed by serial terminal input of network credentials (SSID and password). Real-time pacing parameter adjustments were achieved through IP address-based wireless communication.

### Power System Specifications

The device operated on a 9V battery power source with regulated 5V output through the 7805 voltage regulator. Key voltage specifications included: VBUS (voltage bus) providing 5V output matching battery performance, VSYS serving as the primary power connection point, and 3V3(out) providing secondary 3.3V output for low-power components.

### Outcome Measures

The primary outcome is successful induction of persistent atrial fibrillation, defined as continuous AFib lasting at least 7 days and confirmed by daily 12-lead electrocardiographic monitoring. Secondary outcomes include time to AFib onset (days from initiation of pacing), electrophysiologic remodeling parameters, structural changes, device performance metrics, and safety endpoints. Structural remodeling will be quantified through Masson’s trichrome staining to determine percentage of atrial fibrotic area. Device performance will be evaluated through battery longevity, pacing fidelity (percentage of intended pacing stimuli successfully delivered), and wireless charging efficiency. Safety outcomes will include infection rates, lead dislodgement, ventricular arrhythmias, and mortality.

### Statistical Analysis

The primary endpoint of AFib induction success rate will be analyzed using exact binomial confidence intervals to determine if our observed success rate is consistent with published electrical pacing protocols. Time-to-event data for AFib onset will be visualized using Kaplan-Meier survival curves. For continuous electrophysiologic variables we will use paired t-tests to compare baseline versus post-pacing values within each group, and independent t-tests to compare changes between paced and control groups. Non-normally distributed data, including histological fibrosis scores, will be analyzed using Mann-Whitney U tests for between-group comparisons. All statistical tests will be two-tailed with significance set at p<0.05. Analyses will be performed using SigmaPlot version 16.

All study data is hosted at the University of Arizona and stored in password-protected databases on secure university servers with automated daily backups. Raw data files including ECG recordings, echocardiographic images, electrophysiology tracings, and histological images will be archived in their original formats with unique study identifiers.

### Safety Monitoring and Humane Endpoints

Animal welfare will be monitored through daily clinical assessments including evaluation of appetite, activity level, respiratory effort, and surgical site appearance. Body weight will be measured twice weekly to detect significant weight loss. To minimize risks associated with chronic pacing, pacing output will be limited to less than 10 volts to reduce tissue heating, and daily device interrogation will be performed to detect any malfunctions. Emergency cardioversion equipment will be maintained readily available throughout the study period.

Infection prevention measures include strict sterile surgical technique, and daily surgical site inspection. If infection is suspected based on local erythema, swelling, or drainage, or systemic signs including fever or lethargy, appropriate cultures will be obtained and antimicrobial therapy initiated, with consideration for early device removal if infection is confirmed.

### Study Timeline and Current Status

This protocol is currently in the preparatory phase following successful completion of device validation through pilot testing (described in the Pilot and Preliminary Data section below). The anticipated study timeline spans 12 months from initiation. Months 1-2 will focus on device refinement, specifically completion of the electromagnetic induction charging system for transcutaneous battery recharging and optimization of the charging collar design. Month 3 will involve animal procurement from certified vendors and completion of baseline assessments including electrocardiography, echocardiography, and baseline electrophysiology studies. Months 4-7 will encompass the surgical implantation procedures and chronic pacing protocols. Month 8 will complete all terminal electrophysiology studies. Months 9-10 will be dedicated to histological processing, microscopic analysis, and comprehensive data analysis. Months 11-12 will focus on manuscript preparation and dissemination of results.

### Data Availability and Open Science Commitment

Upon study completion and manuscript publication, all de-identified datasets will be made publicly available in compliance with PLOS One data sharing policies and NIH data sharing guidelines. Datasets to be shared will include raw electrocardiographic data files in standard formats, electrophysiology measurements in spreadsheet format, quantified histological data including fibrosis percentages and cell counts, and device performance metrics.

### Animal Model and In-Vivo Validation Subject Selection and Preparation

Device validation was performed using a domestic swine model with induced myocardial infarction and cardiogenic shock, providing a clinically relevant testing environment that simulates the cardiovascular conditions typical of AFib research subjects.

### Surgical Procedure and Lead Placement

Vascular access was established through the left jugular vein using standard sterile technique. Fluoroscopic guidance was employed throughout the procedure to ensure precise placement of an active fixation pacing lead (Medtronic 5076, Medtronic Inc., Minneapolis, MN). Lead positioning was confirmed through real-time imaging, and electrical parameters were verified prior to connection to the custom pacing device (**Figure 2**).

**Figure 2.**
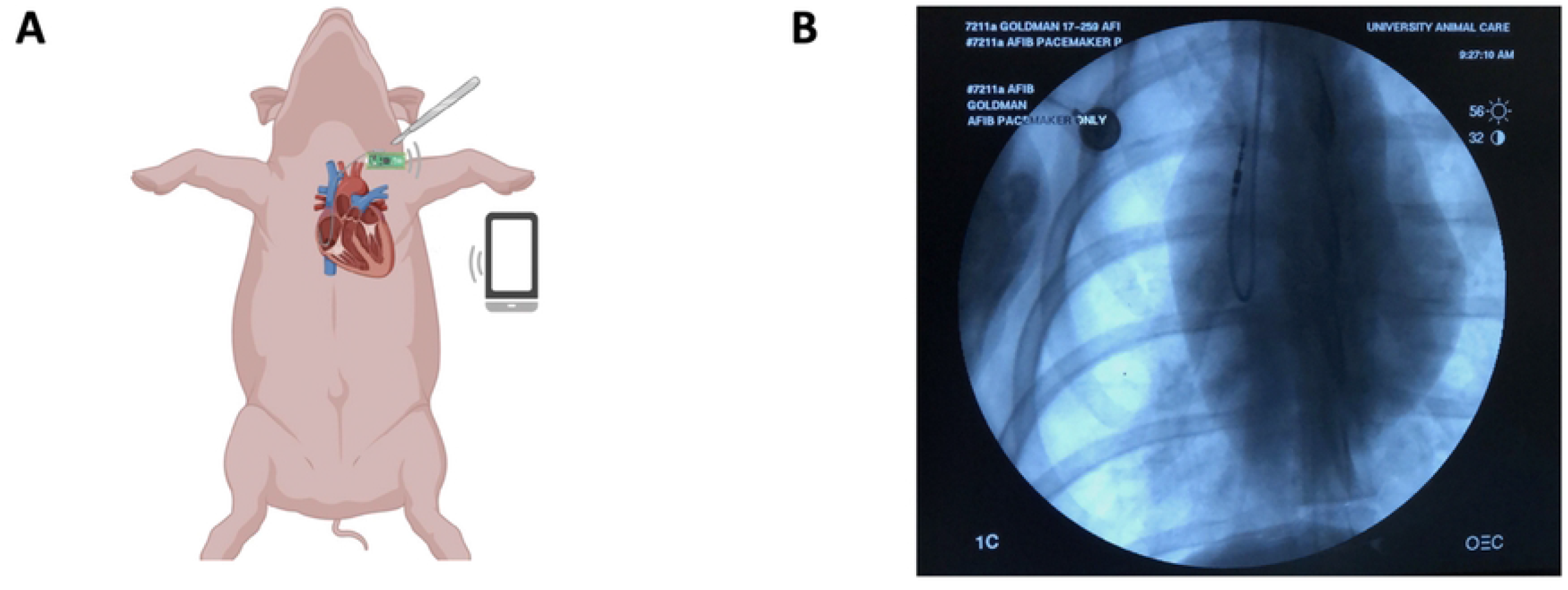
(A) Raspberry Pi Pico W connected to an atrial pacing lead implanted in the right atrium. The device is controlled using a cell phone (B) Active fixation pacing lead (Medtronic 5076) implanted in the right atrium. Created in BioRender. Bravo, F. (2025) https://BioRender.com/rqy7laa

### Electrophysiological Validation

Successful cardiac pacing and capture were confirmed using standard 12-lead electrocardiography. Both atrial and ventricular pacing capabilities were validated to ensure device versatility for future research applications. Pacing thresholds, sensing parameters, and capture confirmation were documented according to standard electrophysiology protocols.

### Pilot and Preliminary Data

To demonstrate the feasibility of our proposed study protocol, we conducted preliminary validation testing of the custom pacing device on an oscilloscope and one domestic swine. The following pilot data supports the technical feasibility of using this device for chronic atrial pacing studies to induce persistent AFib.

### Technical Validation and Electrical Characteristics

Oscilloscope testing was performed and demonstrated frequency and voltage output concordant with the programmed rates. The device demonstrated precise frequency control across the programmable range of 2 Hz to 1.2 kHz. The device consistently maintained variable pacing rates while real-time adjustments were made through the Wi-Fi interface. Additionally, our power supply showed stability throughout pacing sessions, with no malfunctions or interruptions.

The Wi-Fi connectivity of the Raspberry Pi Pico W enabled remote control of pacing parameters without requiring physical access to the device. Real-time frequency adjustments were successfully performed through a web interface, allowing modification of pacing rate during active experiments.

### In-Vivo Cardiac Pacing Validation

The device was tested on a domestic swine. Access was obtained through the left jugular vein and fluoroscopy was used to guide implantation of an active fixation pacing lead (Medtronic 5076). This atrial pacing lead was placed in the right ventricle then relocated to the right atrium with successful pacing with capture was verified using electrocardiography (**Figure 3**). Both atrial and ventricular pacing demonstrated consistent capture thresholds and reliable electrical stimulation, confirming successful tissue interface and electrical delivery.

**Figure 3.**
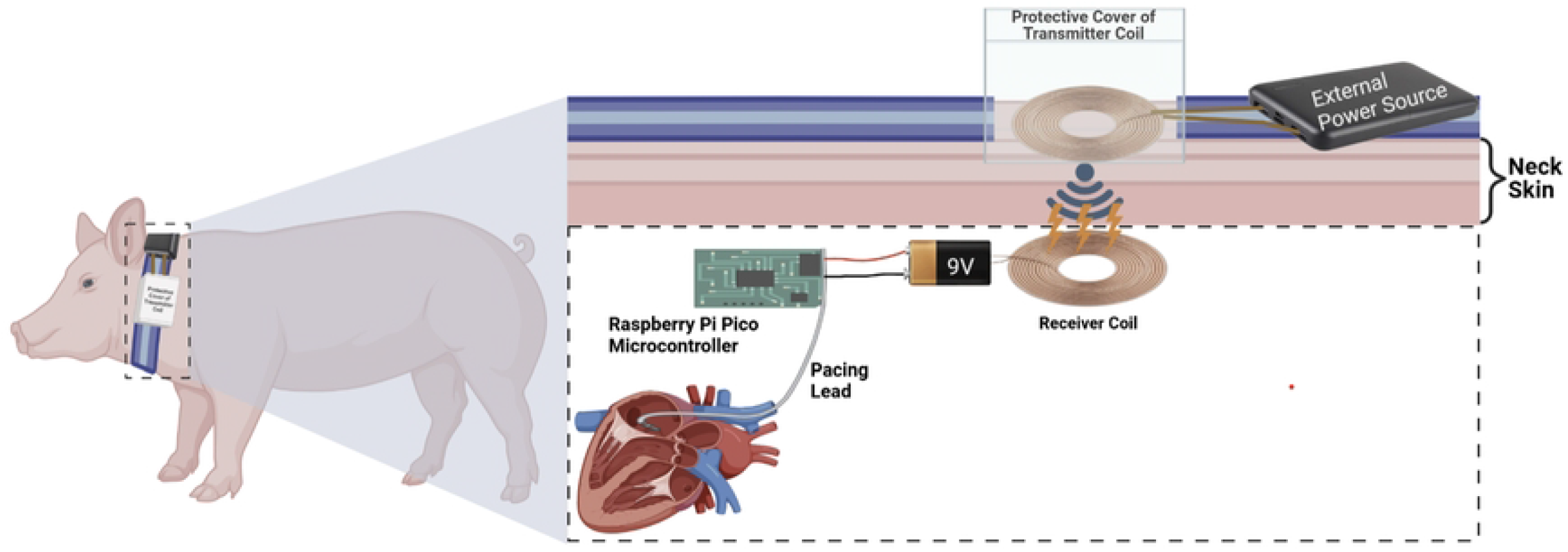
(A) ECG tracing of Atrial Pacing with variable atrioventricular conduction (B) ECG tracing of Ventricular Pacing with capture

### Power System Performance and Reliability

The 9V battery power system with 7805 voltage regulation maintained stable 5V output throughout extended testing sessions. The voltage regulator provides consistent power conversion from the 9V input to the required 5V VSYS connection. No voltage fluctuations or power interruptions were observed during the validation period, confirming the reliability of the power management system for the intended research applications. The modular design allows for separate battery housing to facilitate easier implantation procedures.

## Discussion

Swine models provide multifactorial benefits over rodents, rabbits, dogs, and organoid models to study AFib. Rodents provide genetic modification feasibility (33) but have significant electrophysiologic differences from humans (34). Rabbits and dogs are useful for acute AFib studies (35, 36), however both models have small atria, which restricts the ability to test new ablation or other catheter-based techniques. Previous rabbit studies have been paramount in understanding autonomic effects on AFib (37). Organoids provide high-throughput drug screening and potential for progression towards individualized medicine (38), though they lack systemic influences such as autonomic regulation and hemodynamics, which are known contributors to AFib. Swine provide a good balance between physiological relevance and experimental feasibility, particularly for studying persistent AFib.

Pooled analysis confirms atrial tachypacing effectiveness with ∼99% AFib induction rate. These models have been valuable for testing therapeutics, demonstrating efficacy of novel antiarrhythmics (15). Genetic models have proven useful in studying AFib, with significantly prolonged induction times in modified swine. Future studies may combine techniques (e.g., pacing plus inflammation) to better mimic complex clinical AFib scenarios (30, 39). Swine provide the best compromise between physiological relevance and experimental feasibility, particularly for long-term persistent AFib research.

## Conclusion

Our laboratory study protocol describes a comprehensive plan to validate this low-cost, customizable pacing system, capable of capturing atrial and ventricular tissue. This innovation provides a potentially scalable and efficient alternative method for producing and studying atrial fibrillation pathogenesis and testing new therapies. Having achieved successful pacing in functional assessment, the next step is to use rapid atrial pacing in the right atrium for few weeks in mini swine to induce persistent AFib. This approach yields a large animal model of persistent AFib created using a Raspberry Pi Pico W microcontroller connected to a right atrial pacing lead. Our device offers improved availability and ease of use compared to existing systems. A current limitation of the device is that the 9V battery has a limited lifespan, because once implanted, this device needs to run on the order of weeks. Ideally, a rechargeable battery would be the ideal power source for our device. To solve this problem, we are currently developing an electromagnetic induction charging system that will recharge the battery through the skin. The charging device will be worn by the pig on a collar (**Figure 4**).

**Figure 4.**
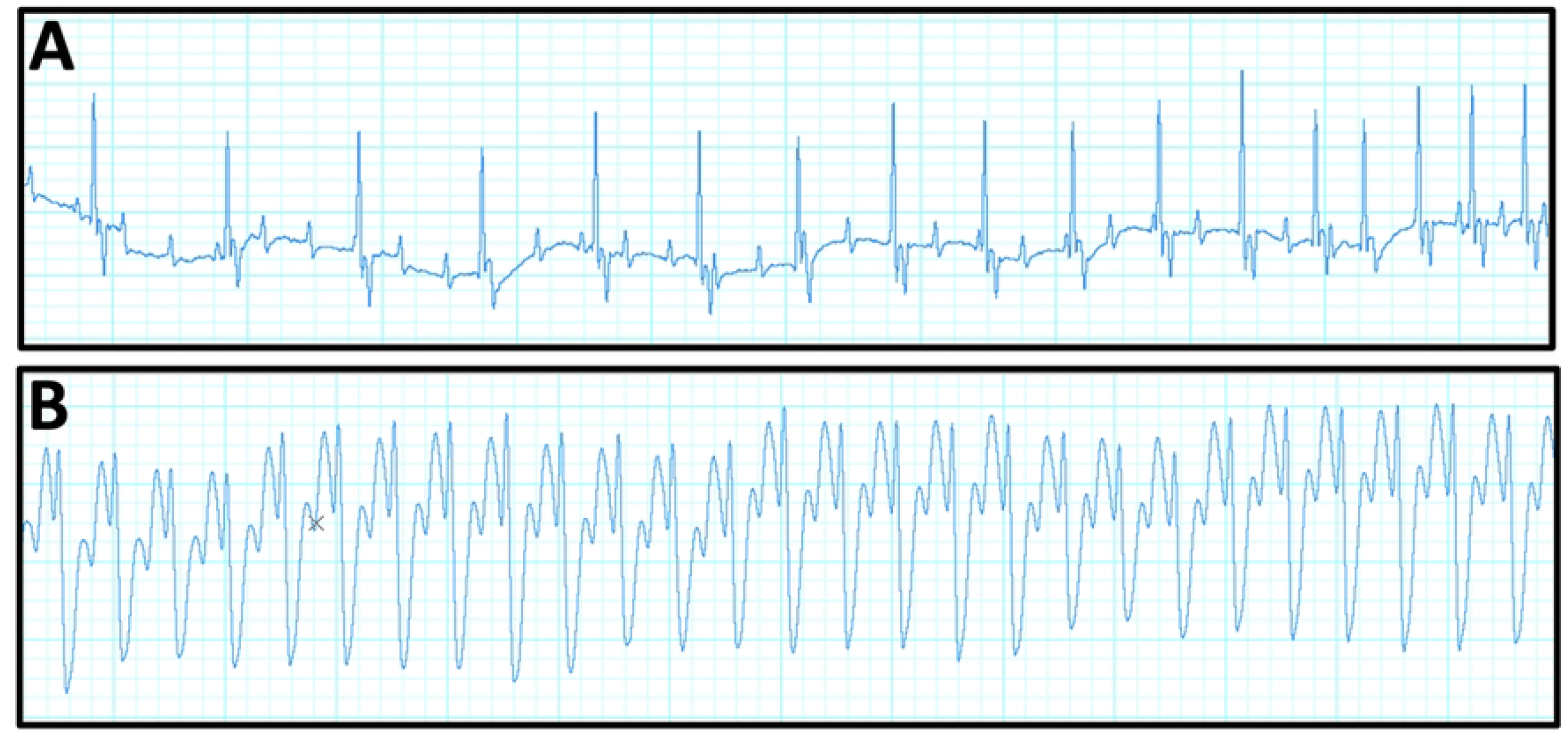
Swine model equipped with electromagnetic induction charging collar and atrial fibrillation induction system. An external transmitter coil transfers voltage wirelessly to a receiver coil subcutaneously, powering the 9V implanted battery to power the Raspberry Pi Pico and pacing lead. Created in BioRender. Bravo, F. (2025) https://BioRender.com/kpcwf5x

The current literature reveals electrical atrial tachypacing most reliably produces persistent AFib. Genetic modifications were found to be effective for producing swine that are more resistant to developing AFib. Pharmacologic and surgical techniques were more useful for inducing short episodes of AFib on the order of minutes and are useful for studying conditions such as post operative AFib. In conclusion, methods of inducing AFib should be tailored whether the acute or chronic process is of study interest. The summary figures highlight methods and their associated timelines currently available to researchers (**Figures 5 and 6**).

**Figure 5.**
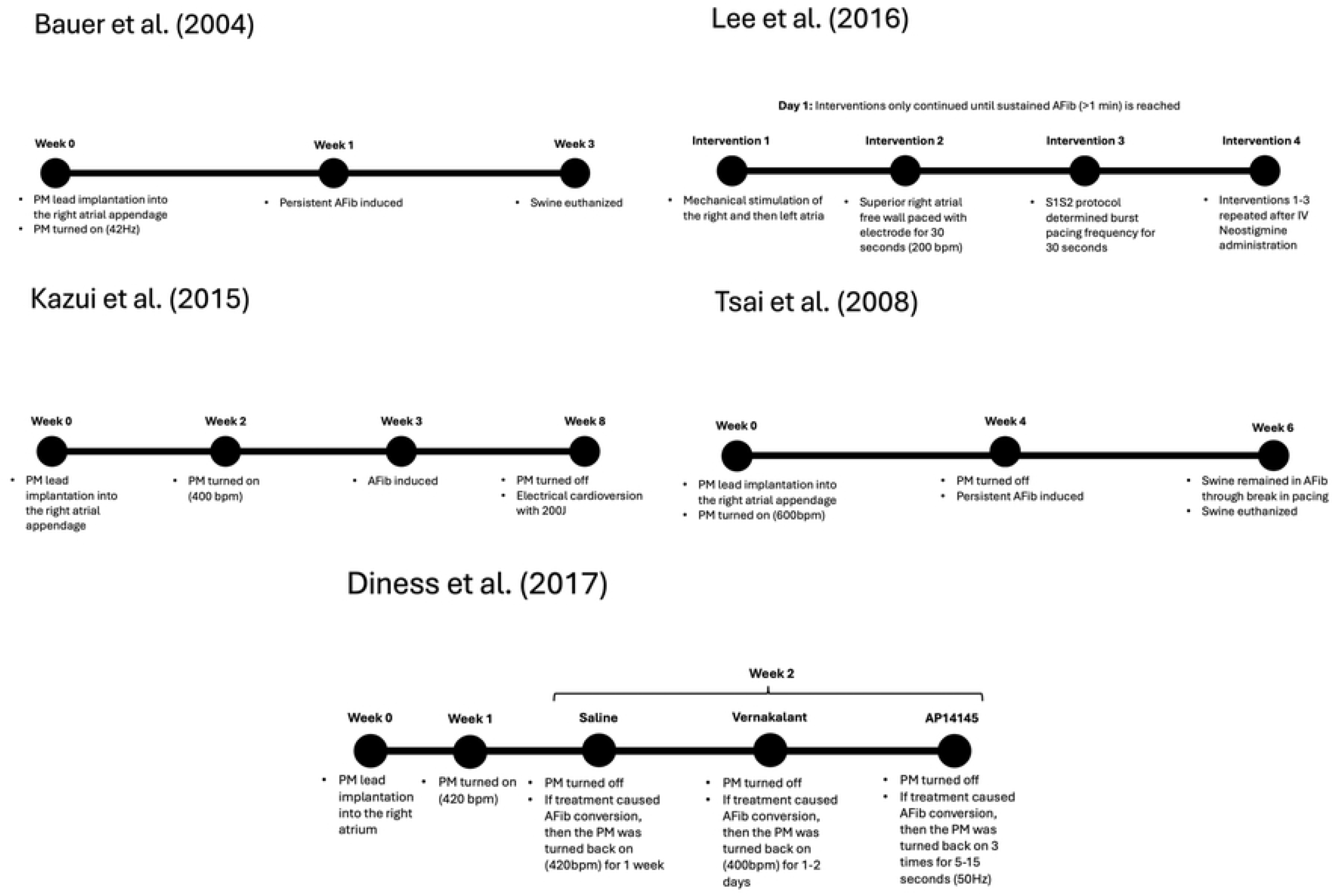
Time points in each of the electrical methods used to produce persistent AFib in swine. Pacemaker is abbreviated here as PM. Exact times for induction and duration of AFib can be seen in Table 1, here we generalize timepoints in terms of weeks.

**Figure 6.**
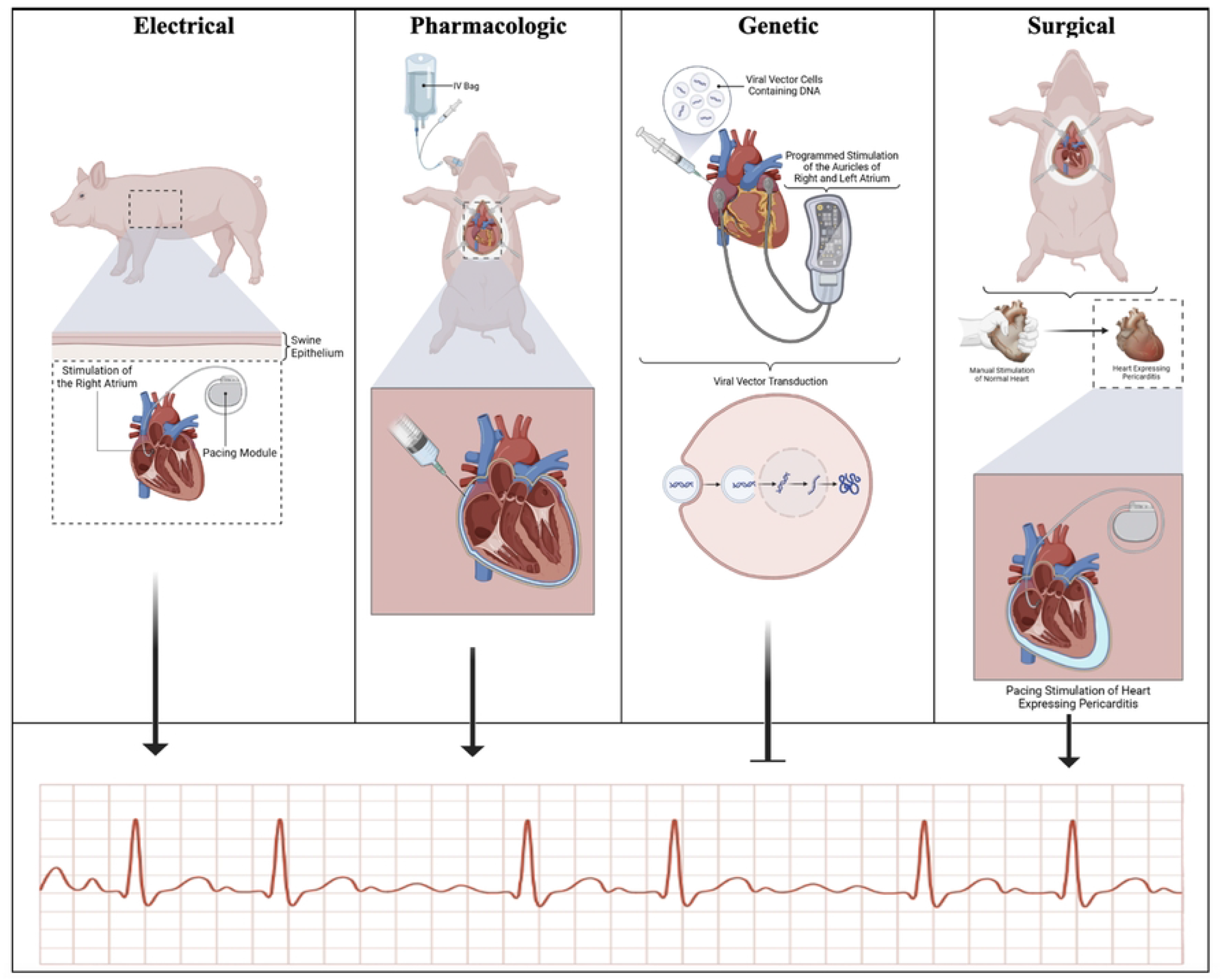
Visual representation of each of the methods reviewed for inducing AFib in swine: electrical, pharmacologic, genetic, and surgical. The figures point to an electrocardiogram displaying AFib and whether they promote AFib induction or inhibit it. The electrical, pharmacologic, and surgical methods stimulate AFib, while genetic models inhibit AFib induction in swine. Created in BioRender. Bravo, F. (2025) https://BioRender.com/8vo6435

## Acknowledgements

A special thank you to everyone in the Goldman lab for their support.

## Contributions

FB – Model design, engineering, experimentation, literature review and writing

JR – Manuscript editing and writing

JR – Figures and review

EL – Figures and manuscript review

PM – Manuscript review and guidance on animal models of AFib

SD – Tables

AG – Technical Support

DB – Technical Support

AST – Manuscript review

MKP – Provided oversight and guidance, edited manuscript.

SG – senior physician-scientist, provided oversight and guidance, assisted with data analysis, edited manuscript.

TM – senior physician-scientist, provided oversight and guidance, assisted with data analysis, edited manuscript.

## References

1. Go AS, Hylek EM, Phillips KA, Chang Y, Henault LE, Selby JV, et al. Prevalence of diagnosed atrial fibrillation in adults: national implications for rhythm management and stroke prevention: the AnTicoagulation and Risk Factors in Atrial Fibrillation (ATRIA) Study. JAMA. 2001;285(18):2370–5.

2. Miyasaka Y, Barnes ME, Gersh BJ, Cha SS, Bailey KR, Abhayaratna WP, et al. Secular trends in incidence of atrial fibrillation in Olmsted County, Minnesota, 1980 to 2000, and implications on the projections for future prevalence. Circulation. 2006;114(2):119–25.

3. Nattel S. Atrial electrophysiological remodeling caused by rapid atrial activation: underlying mechanisms and clinical relevance to atrial fibrillation. Cardiovasc Res. 1999;42(2):298–308.

4. Dzeshka MS, Lip GYH, Snezhitskiy V, Shantsila E. Cardiac Fibrosis in Patients With Atrial Fibrillation: Mechanisms and Clinical Implications. J Am Coll Cardiol. 2015;66(8):943–59.

5. Chen P-S, Chen LS, Fishbein MC, Lin S-F, Nattel S. Role of the Autonomic Nervous System in Atrial Fibrillation. Circ Res. 2014;114(9):1500–15.

6. Roselli C, Rienstra M, Ellinor PT. Genetics of Atrial Fibrillation in 2020. Circ Res. 2020;127(1):21–33.

7. Babapoor-Farrokhran S, Gill D, Rasekhi RT. The role of long noncoding RNAs in atrial fibrillation. Heart Rhythm. 2020;17(6):1043–9.

8. Haïssaguerre M, Jaïs P, Shah DC, Takahashi A, Hocini M, Quiniou G, et al. Spontaneous Initiation of Atrial Fibrillation by Ectopic Beats Originating in the Pulmonary Veins. New England Journal of Medicine. 1998;339(10):659–66.

9. Lin J-L, Lai L-P, Lin C-S, Du C-C, Wu T-J, Chen S-P, et al. Electrophysiological Mapping and Histological Examinations of the Swine Atrium with Sustained (≥24 h) Atrial Fibrillation: A Suitable Animal Model for Studying Human Atrial Fibrillation. Cardiology. 2003;99(2):78–84.

10. Quintanilla JG, Alfonso-Almazán JM, Pérez-Castellano N, Pandit SV, Jalife J, Pérez-Villacastín J, et al. Instantaneous Amplitude and Frequency Modulations Detect the Footprint of Rotational Activity and Reveal Stable Driver Regions as Targets for Persistent Atrial Fibrillation Ablation. Circ Res. 2019;125(6):609–27.

11. Lee AM, Miller JR, Voeller RK, Zierer A, Lall SC, Schuessler RB, et al. A Simple Porcine Model of Inducible Sustained Atrial Fibrillation. Innovations (Phila). 2016;11(1):76–8.

12. Bauer A, McDonald AD, Donahue JK. Pathophysiological findings in a model of persistent atrial fibrillation and severe congestive heart failure. Cardiovasc Res. 2004;61(4):764–70.

13. Tsai CT, Lai LP, Hwang JJ, Chen WP, Chiang FT, Hsu KL, et al. Renin-angiotensin system component expression in the HL-1 atrial cell line and in a pig model of atrial fibrillation. J Hypertens. 2008;26(3):570–82.

14. Citerni C, Kirchhoff J, Olsen LH, Sattler SM, Gentilini F, Forni M, et al. Characterization of Atrial and Ventricular Structural Remodeling in a Porcine Model of Atrial Fibrillation Induced by Atrial Tachypacing. Frontiers in Veterinary Science. 2020;7.

15. Diness JG, Skibsbye L, Simó-Vicens R, Santos JL, Lundegaard P, Citerni C, et al. Termination of Vernakalant-Resistant Atrial Fibrillation by Inhibition of Small-Conductance Ca^2+^-Activated K^+^ Channels in Pigs. Circulation: Arrhythmia and Electrophysiology. 2017;10(10):e005125.

16. Kazui T, Henn MC, Watanabe Y, Kovács SJ, Lawrance CP, Greenberg JW, et al. The impact of 6 weeks of atrial fibrillation on left atrial and ventricular structure and function. J Thorac Cardiovasc Surg. 2015;150(6):1602–8.

17. Diness JG, Kirchhoff JE, Speerschneider T, Abildgaard L, Edvardsson N, Sørensen US, et al. The K(Ca)2 Channel Inhibitor AP30663 Selectively Increases Atrial Refractoriness, Converts Vernakalant-Resistant Atrial Fibrillation and Prevents Its Reinduction in Conscious Pigs. Front Pharmacol. 2020;11:159.

18. Nishida K, Michael G, Dobrev D, Nattel S. Animal models for atrial fibrillation: clinical insights and scientific opportunities. EP Europace. 2009;12(2):160–72.

19. Liu Z, Hutt JA, Rajeshkumar B, Azuma Y, Duan KL, Donahue JK. Preclinical efficacy and safety of KCNH2-G628S gene therapy for postoperative atrial fibrillation. J Thorac Cardiovasc Surg. 2017;154(5):1644-51.e8.

20. Soucek R, Thomas D, Kelemen K, Bikou O, Seyler C, Voss F, et al. Genetic suppression of atrial fibrillation using a dominant-negative ether-a-go-go–related gene mutant. Heart Rhythm. 2012;9(2):265–72.

21. Igarashi T, Finet JE, Takeuchi A, Fujino Y, Strom M, Greener ID, et al. Connexin gene transfer preserves conduction velocity and prevents atrial fibrillation. Circulation. 2012;125(2):216–25.

22. Bikou O, Thomas D, Trappe K, Lugenbiel P, Kelemen K, Koch M, et al. Connexin 43 gene therapy prevents persistent atrial fibrillation in a porcine model. Cardiovasc Res. 2011;92(2):218–25.

23. Park DS, Cerrone M, Morley G, Vasquez C, Fowler S, Liu N, et al. Genetically engineered SCN5A mutant pig hearts exhibit conduction defects and arrhythmias. J Clin Invest. 2015;125(1):403–12.

24. Carneiro JS, Bento AS, Bacic D, Nearing BD, Rajamani S, Belardinelli L, et al. The Selective Cardiac Late Sodium Current Inhibitor GS-458967 Suppresses Autonomically Triggered Atrial Fibrillation in an Intact Porcine Model. J Cardiovasc Electrophysiol. 2015;26(12):1364–9.

25. Fuller H, Justo F, Nearing BD, Kahlig KM, Rajamani S, Belardinelli L, et al. Eleclazine, a new selective cardiac late sodium current inhibitor, confers concurrent protection against autonomically induced atrial premature beats, repolarization alternans and heterogeneity, and atrial fibrillation in an intact porcine model. Heart Rhythm. 2016;13(8):1679–86.

26. Justo F, Fuller H, Nearing BD, Rajamani S, Belardinelli L, Verrier RL. Inhibition of the cardiac late sodium current with eleclazine protects against ischemia-induced vulnerability to atrial fibrillation and reduces atrial and ventricular repolarization abnormalities in the absence and presence of concurrent adrenergic stimulation. Heart Rhythm. 2016;13(9):1860–7.

27. Schwarzl M, Alogna A, Zweiker D, Verderber J, Huber S, Manninger M, et al. A porcine model of early atrial fibrillation using a custom-built, radio transmission-controlled pacemaker. J Electrocardiol. 2016;49(2):124–31.

28. Li B, Cui Y, Zhang D, Luo X, Luo F, Li B, et al. The characteristics of a porcine mitral regurgitation model. Exp Anim. 2018;67(4):463–77.

29. Shi Y, Zhang H, Huang S, Yin L, Wang F, Luo P, et al. Epigenetic regulation in cardiovascular disease: mechanisms and advances in clinical trials. Signal Transduction and Targeted Therapy. 2022;7(1):200.

30. Clauss S, Schüttler D, Bleyer C, Vlcek J, Shakarami M, Tomsits P, et al. Characterization of a porcine model of atrial arrhythmogenicity in the context of ischaemic heart failure. PLoS One. 2020;15(5):e0232374.

31. Lee S, Khrestian C, Laurita D, Juzbasich D, Wallick D, Waldo A. Validation of a new species for studying postoperative atrial fibrillation: Swine sterile pericarditis model. Pacing Clin Electrophysiol. 2023;46(8):1003–9.

32. Kumagai K, Khrestian C, Waldo AL. Simultaneous Multisite Mapping Studies During Induced Atrial Fibrillation in the Sterile Pericarditis Model. Circulation. 1997;95(2):511–21.

33. Ozcan C, Battaglia E, Young R, Suzuki G. LKB1 Knockout Mouse Develops Spontaneous Atrial Fibrillation and Provides Mechanistic Insights Into Human Disease Process. Journal of the American Heart Association. 2015;4(3):e001733.

34. Ferreira M, Geraldes V, Felix AC, Oliveira M, Laranjo S, Rocha I. Advancing Atrial Fibrillation Research: The Role of Animal Models, Emerging Technologies and Translational Challenges. Biomedicines. 2025;13(2):307.

35. Ravens U, Peyronnet R. Electrical Remodelling in Cardiac Disease. Cells. 2023;12(2):230.

36. Odening KE, Gomez A-M, Dobrev D, Fabritz L, Heinzel FR, Mangoni ME, et al. ESC working group on cardiac cellular electrophysiology position paper: relevance, opportunities, and limitations of experimental models for cardiac electrophysiology research. EP Europace. 2021;23(11):1795–814.

37. Oliveira M, Postolache G, Geraldes V, Silva V, Laranjo S, Tavares C, et al. Modulação eletrofisiológica das aurículas e veias pulmonares: interação da estimulação simpática e parassimpática na inducibilidade de fibrilhação aguda auricular num modelo experimental in vivo. Rev Port Cardiol. 2012;31(3):215–23.

38. Cofiño-Fabres C, Passier R, Schwach V. Towards Improved Human In Vitro Models for Cardiac Arrhythmia: Disease Mechanisms, Treatment, and Models of Atrial Fibrillation. Biomedicines. 2023;11(9):2355.

39. Schüttler D, Bapat A, Kääb S, Lee K, Tomsits P, Clauss S, et al. Animal Models of Atrial Fibrillation. Circ Res. 2020;127(1):91–110.

